# Cholesterol Metabolite 27-Hydroxycholesterol Enhances the Secretion of Cancer Promoting Extracellular Vesicles by a Mitochondrial ROS-Induced Impairment of Lysosomal Function

**DOI:** 10.1101/2024.05.01.591500

**Authors:** Anasuya Das Gupta, Jaena Park, Janet E. Sorrells, Hannah Kim, Natalia Krawczynska, Hashni Epa Vidana Gamage, Adam T. Nelczyk, Stephen A. Boppart, Marni D. Boppart, Erik R. Nelson

## Abstract

Extracellular vesicles (EVs) serve as crucial mediators of cell-to-cell communication in normal physiology as well as in diseased states, and have been largely studied in regard to their role in cancer progression. However, the mechanisms by which their biogenesis and secretion are regulated by metabolic or endocrine factors remain unknown. Here, we delineate a mechanism by which EV secretion is regulated by a cholesterol metabolite, 27-Hydroxycholesterol (27HC), where treatment of myeloid immune cells (RAW 264.7 and J774A.1) with 27HC impairs lysosomal homeostasis, leading to shunting of multivesicular bodies (MVBs) away from lysosomal degradation, towards secretion as EVs. This impairment of lysosomal function is caused by mitochondrial dysfunction and subsequent increase in reactive oxygen species (ROS). Interestingly, cotreatment with a mitochondria-targeted antioxidant rescued the lysosomal impairment and attenuated the 27HC-mediated increase in EV secretion. Overall, our findings establish how a cholesterol metabolite regulates EV secretion and paves the way for the development of strategies to regulate cancer progression by controlling EV secretion.

## INTRODUCTION

Extracellular vesicles (EVs) are nano-sized membrane bound vesicles comprised of biologically active cargo including proteins, lipids and nucleic acids (1,2). They are signaling entities involved in mediating cross talk between cells and in this way behave like multi-signaling hormones (3). The International Society for Extracellular Vesicles (ISEV) now defines EVs as “particles naturally released from the cell that are delimited by a lipid bilayer and cannot replicate” (4). EVs are broadly divided into two subgroups: small EVs or exosomes, which fall in the size range of 30-150 nm and are derived from endosomes by their maturation to form multivesicular bodies (MVBs), and large EVs or microvesicles (MVs), which are in the size range of 150-500 nm and are formed at the plasma membrane (5). Other sub-types of EVs include exomeres (6), oncosomes (7) and apoptotic bodies (8).

EVs, in addition to other secreted factors such as chemokines and growth factors, are critical messengers in tumor progression and metastasis (9,10). EVs influence cancer progression by transporting molecules that can initiate several pathophysiological processes such as vascular leakiness, angiogenesis, and formation of the pre-metastatic niche (11,12). Therapeutic targeting of EVs and EV cargo could provide an approach to impair metastasis. Several factors such as post-translational modifications (13,14), hypoxia (15,16), proto-oncogenes and oncogenes (17,18), and metabolic changes such as alterations in rates of glycolysis and oxidative phosphorylation (19,20) have been implicated in modulating EV secretion as well as EV cargo. Targeting of these regulatory pathways may serve as a means to impair the secretion of cancer promoting EVs and thus act as potential therapeutic strategies for cancer. Specifically, pathways that alter EV secretion may be prime targets for such intervention. However, neither the metabolic or endocrine agents that regulate EV secretion, nor the mechanisms by which EV secretion is regulated, are fully understood.

Elevated cholesterol has been identified as a poor prognostic factor for breast cancer patients and cholesterol lowering drugs such as statins have been associated with increased recurrence free survival (21). The cholesterol metabolite, 27-Hydroxycholesterol (27HC), is an endogenous oxysterol, that has been shown to mediate many of the pro-tumor effects of cholesterol (22–24). Physiologically, 27HC acts as a regulator of cholesterol homeostasis, through feedback inhibition of cholesterol biosynthesis, promotion of cholesterol catabolism and through increased cellular cholesterol efflux (25). However, in the context of breast cancer, 27HC has been shown to increase the proliferation of estrogen receptor positive (ER+) cancer cells in an ER dependent manner (22). In addition, 27HC treatment of metastasis-bearing mice led to increased myeloid immune cell infiltration, and a decrease of intratumoral CD8+ cytotoxic T cells (26). Mechanistically, one way in which 27HC promotes breast cancer progression is through its action on liver x receptors (LXRs) in myeloid cells, resulting in suppression of T-cell expansion (24). Collectively, these studies strongly implicate the involvement of 27HC and myeloid immune cells in the progression of breast cancer.

Interestingly, while exploring how 27HC alters myeloid immune cell function, we found that the treatment of several cell types with 27HC resulted in an increased rate of EV secretion and altered EV cargo (27). Importantly, primary tumor and metastatic burden was increased in mice treated with EVs derived from myeloid immune cells incubated with 27HC; findings that were consistent in two different murine mammary cancer models, 4T1 and Met1 (27). Given that EVs secreted post-27HC treatment are pro-tumorigenic, our objective was to elucidate the mechanism by which 27HC increases the biogenesis of these EVs.

Here we have established that 27HC increases EV secretion by impairing lysosomal function, leading to the MVBs being directed away from lysosomal degradation and towards secretion as EVs. We have also observed that the impairment of lysosomal function is caused in part by 27HC-mediated mitochondrial disruption and subsequent production of reactive oxygen species. Further, we have observed that co-treatment of 27HC with Mitoquinol (mitoQ), a mitochondria targeted antioxidant, is able to rescue the lysosomal damage and therefore attenuate the 27HC mediated increase in EV secretion. Overall, we have delineated a mechanism causing the increased secretion of EVs.

## MATERIALS AND METHODS

### Reagents

27HC (purity 95%) was synthesized by Sai Labs (Hyderabad, India). Bafilomycin A and Mitoquinol were purchased from Cayman Chemicals (Ann Arbor, MI). All compounds were dissolved in DMSO and stored at -20°C.

### Cell Culture

Murine cell lines RAW 264.7 and J774A.1 were purchased from the American Type Culture Collection (ATCC, USA) and were cultured in Dulbecco’s Modified Eagle Medium (DMEM, Gibco, USA) media, supplemented with 10% fetal bovine serum (FBS, Gibco), 1% Non-essential amino acids (Corning), 1% Sodium Pyruvate (Corning) and 1% Penicillin/Streptomycin (Corning). Briefly, cells were seeded and treated with the respective compounds for 24 hours in media containing EV-depleted FBS (Gibco). All cells were maintained at 37°C with 5% CO2. Cell lines were cultured no longer that passage 25. Cell lines were routinely tested for mycoplasma.

### EV Isolation

EVs were isolated from cell culture supernatant using one of the two independent methods. For nanoparticle tracking analysis (NTA) EVs were enriched using ExoQuick TC ULTRA EV Isolation Kit (SBI Biosciences) as previously described (27). For NTA and Western blot analyses EVs were enriched by differential centrifugation which collected the supernatants after 800 x g for 10 mins and 2,000 x g for 30 mins followed by ultracentrifugation (Sorvall, Thermo Scientific) at 10,000xg for 1 hour (Microvesicles or large EVs) and 100,000xg for 1 hour (exosomes or small EVs). Supernatant depleted of EVs was considered as vesicle free media and used for western blot analysis.

### Nanoparticle Tracking Analysis (NTA)

The concentration of EVs and their size distributions were measured using NanoSight NS300 (Malvern Panalytical). For each sample, three 30 second videos were captured and analyzed using NTA3.1 software. During capture, screen gain was set at 1.0 and camera level at 13. For analysis, screen gain was set at 10.0 and detection setting at 5.

### Western Blot Analysis

For Western blot analysis of cells, lysis was carried out using RIPA lysis buffer and protein was quantified using BCA Assay (Thermo Fischer Scientific, USA). An equal amount of cell protein was used, and protein was separated using SDS-PAGE. For EVs, EV protein obtained from equal number of cells was lysed directly in Laemmli buffer and loaded into the gel. Separated proteins were transferred to PVDF membrane and blocked using 5% milk in TBST. The blots were incubated with primary antibody overnight at 4°C, washed using TBST, incubated with HRP-conjugated secondary antibody for 1 hour at RT and washed using TBST. Heat shock protein 90 (HSP90) was used as loading control. Blots were incubated with either SuperSignal West Pico or Femto chemiluminescent substrates and imaged using iBrightCL1000 imaging system (Thermo Fischer Scientific, USA).

### RNA isolation and quantitative PCR (qPCR) analysis

Total RNA was extracted from cells using GeneJet RNA Purification kit (Thermo Fischer) and cDNA was synthesized using iScript Reverse Transcription Supermix (Bio-Rad), as previously described (27). Gene expression was determined by qPCR, using iTaq Universal SYBR Green Supermix (Bio-Rad). The primers used were designed using Primer-Blast (https://www.ncbi.nlm.nih.gov/tools/primer-blast/). Expression was determined using the formula, 2^-ΔΔCT^, and normalized to housekeeping gene, TATA box binding protein (TBP).

### Immunofluorescence Microscopy

Cells were cultured in glass slides (ibidi) coated with poly-L-lysine (Sigma) and treated with respective compounds prior to being fixed and permeabilized with ice cold methanol for 5 minutes, blocked with 5% BSA for 1 hour and then incubated with indicated primary antibodies (anti-EEA1, anti-CD63, anti-LAMP1 and anti-CTSB; 1:100; Abcam) at 4°C overnight, followed by secondary antibody (Goat polyclonal Ab to Rabbit IgG (Alexa Fluor®488); 1:1000; Abcam) at room temperature for 1 hour. Samples were mounted with mounting medium containing DAPI (ibidi) and imaged using Ziess LSM700 or LSM900 Confocal Microscope. Images were analyzed using Fiji.

### (Scanning) Transmission Electron Microscopy

For analysis of MVBs using electron microscopy (EM), cells were pelleted down and fixed using 2.0 % paraformaldehyde and 2.5% glutaraldehyde (both E.M. grade) in 0.1 M Na-Cacodylate buffer, pH 7.4, for 4 hours at 4°C. Samples were washed in Na-Cacodylate buffer for 10 minutes and again fixed in 1.0% aqueous osmium tetroxide for 90 minutes in the dark. Samples were washed and then stained using 2% aqueous uranyl acetate, overnight, at 4°C. Then, samples were dehydrated using a graded ethanol series (37%, 67%, 95%) for 10 minutes each and then using 100% ethanol 3 times for 10 minutes each on a shaker. They were then infiltrated using a series of ethanol:polypropylene oxide for 10 minutes each and propylene oxide:Polybed 812 mixture (without DMP-30) for 10 minutes each, prior to embedding. Samples were then embedded in 100% Polybed 812 mixture (without DMP-30), overnight at room temperature. Then they were placed in 100% Polybed 812 mixture with 1.5% DMP-30, in molds, and into oven at 60°C for 24 hours. Thin sections were cut using an ultramicrotome (Ultracut UCT, Leica), collected on grids and examined by EM (Tecnai G2 F20 S-Twin 200kV, Thermo FEI).

### Measurement of Number of acidic particles and Lysosomal pH

Acidic particles in cells were measured using LysoTracker™ Deep Red dye (Invitrogen, USA) and Lysosomal pH was measured using LysoSensor™ Green DND-189 dye (Invitrogen, USA). Cells were cultured in ibidi glass slides and treated with respective compounds. After incubation, the media was replaced with media containing the respective dye, along with Hoesht 33342 dye (Invitrogen, USA) to stain the nucleus. Samples were imaged using Ziess LSM900 Confocal Microscope. Images were analyzed using Fiji. Signal was quantified per field of view and normalized to Hoesht.

### Measurement of Cellular Oxidative Stress and Mitochondrial Function

The amount of reactive oxygen species (ROS) produced within the cell was measured using CellROX™ Deep Red Reagent for oxidative stress detection (Invitrogen, USA). The cells were cultured in glass slides (ibidi) coated with poly-L-lysine (Sigma) and treated with respective compounds. CellROX reagent was added to media, along with Hoesht 33342 dye (Invitrogen, USA). Samples were imaged using Ziess LSM900 Confocal Microscope. Images were analyzed using Fiji. Signal was quantified per field of view and normalized to Hoesht. Mitochondrial mass, Mitochondrial membrane potential, and Mitochondrial ROS were measured similarly using MitoTracker Deep Red, TMRM and MitoSOX dyes.

### Label-free Imaging using Simultaneous Label-free Autofluorescence Multiharmonic (SLAM) Microscopy

Imaging of cells using SLAM Microscopy was carried out as described in (27,28). In short, cells were cultured in glass bottom dish coated with poly-L-lysine and treated with respective compounds. Cells were imaged using multiphoton microscopy with excitation centered at 110 nm with an average power of 14 mW pulse-shaped beam to excite the sample and NAD(P)H and FAD signals were captured using 420-480 nm and 580-640 nm filter, at a spatial resolution of <500nM.

### Seahorse Assay

Cells were seeded in XFe96 cell culture microplates (Agilent, USA) and treated with compounds for 24 hours. Cells were washed with Seahorse XF DMEM assay buffer (Agilent, USA) supplemented with 10mM glucose, 1mM pyruvate and 2mM glutamine, and incubated for 1 hour at 37°C without CO_2_. The ATP production from mitochondrial respiration and glycolytic respiration in response to Oligomycin and Rotenone/Antimycin A was measured using the ATP production assay kit (Agilent, USA). The oxygen consumption rate (OCR) and Extracellular acidification rate (ECAR) from mitochondrial oxidative phosphorylation (OXPHOS) in response to Oligomycin, Carbonyl cyanide-p-trifluoromethoxyphenylhydrazone (FCCP) and Rotenone/Antimycin A was measured using the mitochondrial stress test kit (Agilent, USA). The glycolytic activity or glycoPER in response to Rotenone/Antimycin A and 2-deoxy-D-glucose was measured using glycolytic rate assay (Agilent, USA). All measurements were performed using Seahorse XFe96 Bioanalyzer (Agilent, USA).

### Statistical Analysis

All statistical analysis was performed using GraphPad Prism software. Data are presented as mean+/-SEM. Statistical measurement was carried out using Student’s t-test for two groups and using Ordinary one-way ANOVA with Dunnett’s multiple comparison test for more than two groups. A p-value <0.05 was considered statistically significant.

## RESULTS

### 27-Hydroxycholesterol (27HC) promotes EV secretion in two different myeloid immune cell lines

We have previously shown that 27HC stimulated EV secretion in myeloid immune cells and that the EVs from 27HC-treated myeloid cells had a different size profile. Here, we explored the extent of this regulation by examining its effects in two different cell line models of myeloid cells: RAW 264.7 and J774.A1. The RAW264.7 line is a monocyte/macrophage-like line derived from a tumor induced by the Abelson murine leukemia virus and is a very commonly used mouse ‘macrophage’ line in medical research (28,29). J774.A1 are derived from a female mouse with reticulum cell sarcoma and have monocyte/macrophage morphology and have been used to explore various macrophage functions (30–32). Therefore, we first wanted to explore whether the regulation of EV secretion by 27HC extended to these different myeloid cell models.

Cells were treated with 27HC (10µM) for 24 hours, the EC_50_ of 27HC for either the estrogen receptor or liver x receptor being around 1µM (33,34). EVs were then isolated from the conditioned media using ultracentrifugation, considered the gold standard of EV enrichment. Nanoparticle tracking analysis (NTA) of EV particle number and size distribution indicated that 27HC treated RAW 264.7 cells resulted in increased EV secretion (**Fig. 1A**). Furthermore, the particle size distribution was shifted towards a slightly larger size (**Fig. 1B**). These findings were also found when EVs were isolated using a commercial kit (ExoQuick, **Fig. 1C-D**). Importantly, similar findings were observed when using the J774A.1 model, indicating that 27HC-induced EV secretion is not cell-line specific (**Fig. 1E-H**). We characterized these EVs as expressing the classic EV tetraspanin markers CD63 and CD9, as well as ALIX (**Fig. 1I**). These data are consistent with our previously published findings, where EVs from RAW 264.7 cells were characterized using flow cytometry for tetraspanins and transmission electron microscopy for ultrastructure (35). Therefore, we conclude that RAW264.7 and J774.A1 cell lines are a suitable model to explore the mechanisms by which 27HC stimulates EV biogenesis.

**Figure 1:**
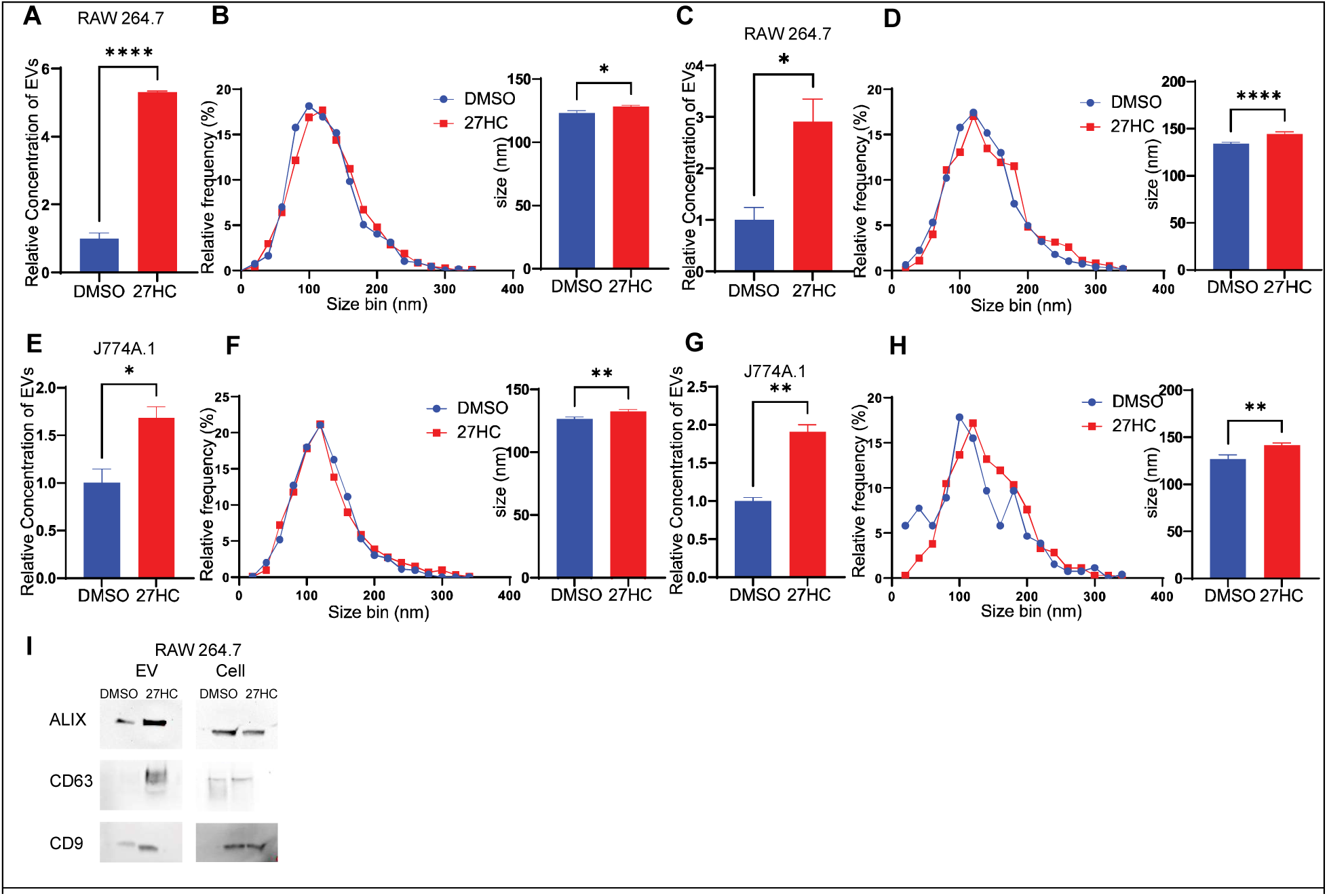
27-Hydroxycholesterol (27HC) promotes EV secretion from cell line models of myeloid immune cells. EVs were collected from conditioned media of RAW 264.7 cells (**A-D**) and J774A.1 cells (**E-H**), using two different EV isolation techniques (Ultracentrifugation and ExoQuick Kit). The number of particles and size distribution were measured by Nanoparticle Tracking Analysis (NTA) using NanoSight NS300 (n=3/condition). (**I**) EVs were characterized for standard EV markers using western blot. EVs from equal number of cells were used for the blots. Statistical analyses were performed using Student’s t-test. ****P-value<0.0001, **P-value<0.01, *P-value<0.05. Data are presented as mean+/-SEM.

### 27-Hydroxycholesterol (27HC) treatment does not alter genes associated with endosome formation, but leads to the enlargement of multivesicular bodies (MVBs)

EVs are a heterogeneous population consisting of exosomes and microvesicles. Although still understudied, the process of EV biogenesis begins with the formation of early endosomes at the plasma membrane, which then form late endosomes. A complex machinery called the Endosomal Sorting Complex Required for Transport (ESCRT) is recruited for the sorting of cargo. The endosome then matures to form multivesicular bodies (MVBs) containing intraluminal vesicles (ILVs) (36,37). These MVBs can either be degraded in the lysosome or, with the help of Rab and SNARE transport proteins, dock at the plasma membrane and release the ILVs as exosomes (3,38–40). Microvesicles are formed by outward budding and fission at the plasma membrane (40–42).

To determine the mechanism by which 27HC modulates EV secretion, we first analyzed the expression of several genes associated with the biogenesis of EVs, expecting that an upregulation of these genes would be required for increased biogenesis. We found that 27HC does not significantly upregulate the expression of genes encoding ESCRT components (**Fig. 2A**), in either RAW264.7 or J774A.1 cells. This indicates that either the modulation of ESCRT is not at the mRNA level, or does not involve ESCRT in its regulation of EV biogenesis.

**Figure 2:**
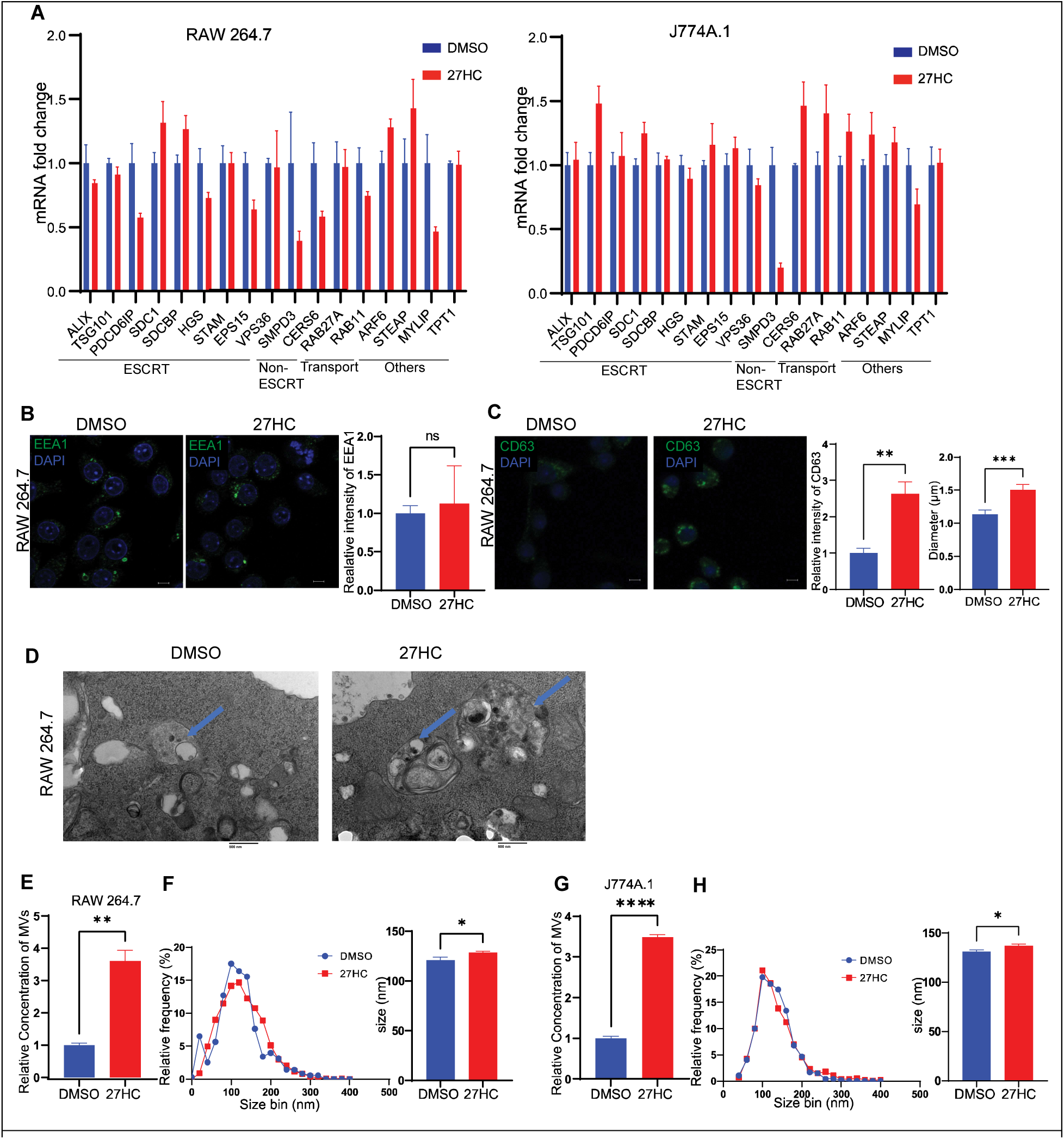
27HC treatment does not alter genes associated with endosome formation, but leads to the enlargement of multivesicular bodies (MVBs). (**A**) mRNA expression of several ESCRT-related, non-ESCRT-related and other genes associated with EV biogenesis indicates that 27HC does not lead to increased expression. RAW 264.7 or J774A.1 cells were treated with DMSO (control) or 27HC for 24hrs prior to assessment of mRNA by RT-qPCR. (**B**) Confocal microscopy analysis of Early Endosomal marker, EEA1 (green) indicating that 27HC does not alter the amount of early endosomes. Scale bar, 5um. Right graph: quantification of EEA1 signal relative to DAPI (n=4). Each data point corresponds to the intensity of EEA1 signal per field of view. (**C**) Confocal microscopy analysis of MVB marker, CD63 (green) indicating that 27HC increases the amount of CD63 intensity per cell and increases the size of CD63 positive MVBs in the cells. Scale bar, 5um. Right graphs: quantification of CD63 signal relative to DAPI (n=3) and size of CD63 positive MVBs (n=20 dots). Each data point corresponds to the intensity of CD63 per field of view and size of CD63 positive MVBs. (**D**) (S)TEM of cells treated with DMSO (control) or 27HC indicating enlarged MVBs (blue arrows) Scale bar, 100nm. (**E-H**) EVs obtained from the 10k ultracentrifugation fraction indicating and increase in number of particles and size distribution in RAW 264.7 cells and J774A.1 cells, on treatment with 27HC. Statistical analyses were performed using a Student’s t-test. ****P-value<0.0001, ***P-value<0.001, **P-value<0.01, *P-value<0.05. Data are presented as mean+/-SEM. Original confocal images were used for analysis and representative images depicted here were brightness adjusted.

Therefore, we next focused on biogenesis mechanisms downstream of the ESCRT pathway. Immunofluorescence (IF) analysis of the endosome marker EEA1 indicated that 27HC did not change the total intensity or the size of EEA1-positive early endosomes, suggesting that the regulation is not at the endosomal level (**Fig. 2B**). However, 27HC did increase the levels of the MVB marker CD63 and increased the size of the CD63-positive MVBs, as assessed by IF (**Fig 2C**). Transmission electron microscopy also revealed an enlargement in the size of vesicular bodies within the cells (**Fig. 2D**). Interestingly, when the MV fraction of ultracentrifuged conditioned media was assessed, 27HC was found to also increase the presence and size of these larger particles (**Fig. 2E-H**). Collectively, these data suggest that the mechanism by which 27HC increases EV secretion (1) is likely to impact the secretion of both small and large EVs, and (2) be downstream of the ESCRT complex.

### 27-Hydroxycholesterol impairs lysosomal function

Thus far, our data indicates that 27HC increases the secretion of both small and large EVs and seems to work downstream of the ESCRT complex. This points to a mechanism whereby treatment with 27HC leads to the secretion of EVs that would otherwise have been targeted towards lysosomal degradation. We have observed that 27HC increases MVB size (**Fig. 2C&D**). This indicates that there is less degradation and recycling of the MVBs within the cell and instead, the MVB cargo is being released as EVs. Interestingly, there have been several reports that an increase in MVB size can result from decreased trafficking of MVBs towards the lysosome (13,43). Lysosomes are a crucial regulatory component of the cell and is involved in maintaining cellular homeostasis by actively degrading and recycling intracellular components and is also involved in regulating cellular signaling and metabolism (44,45). Lysosomes act as the fate determining step for MVBs once they are formed-MVBs can either fuse with the lysosome and get degraded, where the cargo gets recycled within the cell, or the MVBs can fuse with the plasma membrane and release its contents as EVs (46). To study the effect of 27HC on lysosomes, we stained for LAMP1, which is a lysosomal marker, and observed no significant decrease in the overall LAMP1 signal intensity in the cell (**Fig. 3A**). However, there was an increase in the size of those lysosomes remaining after 27HC-treatment (**Fig. 3A**).

**Figure 3:**
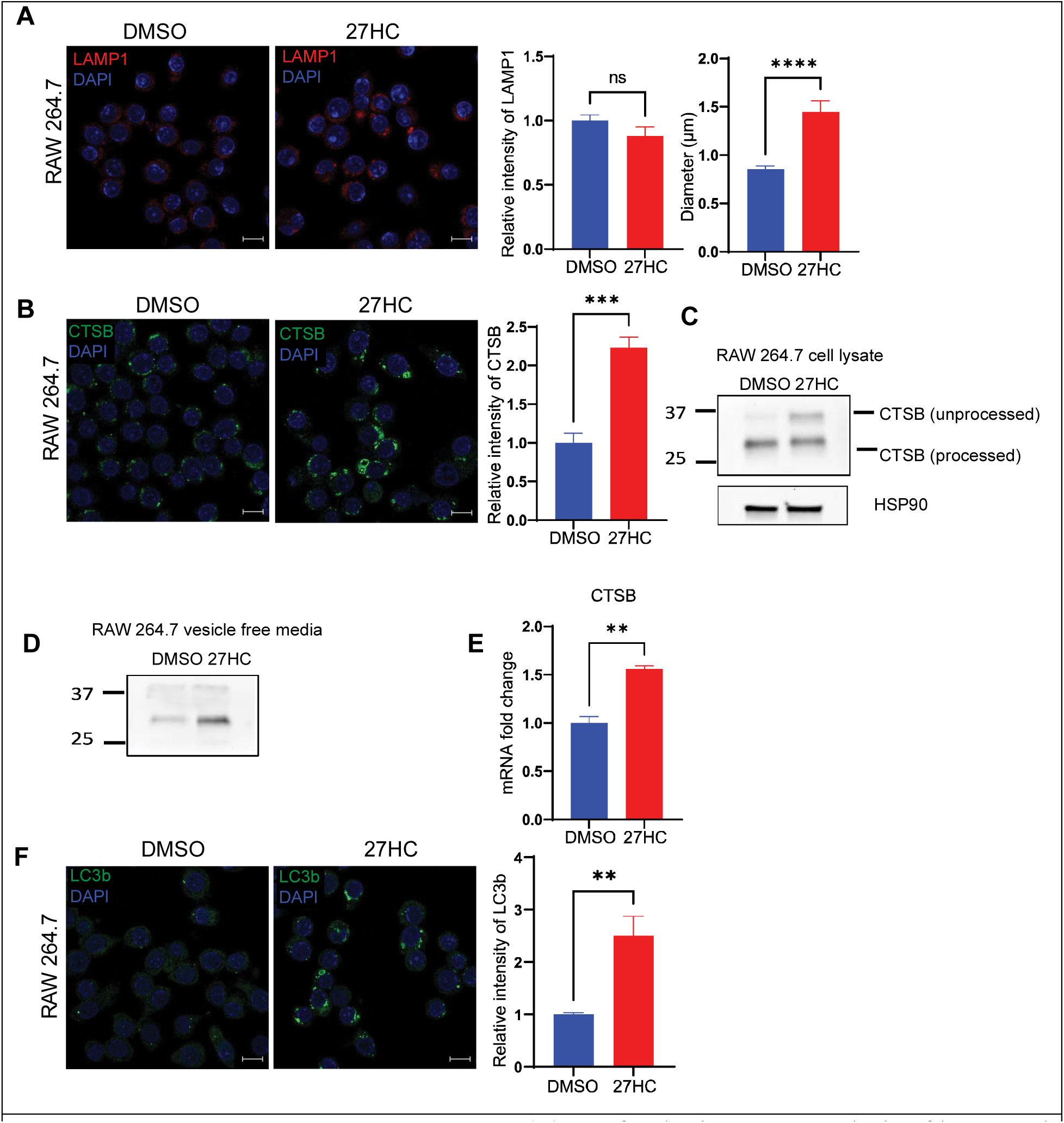
27HC impairs lysosomal function. (**A**) Confocal microscopy analysis of lysosomal marker, LAMP1 (red) indicating that 27HC increases the size of LAMP1 positive lysosomes in the cells. Scale bar, 5um. Right graphs: quantification of LAMP1 signal relative to DAPI (n=5) and size of LAMP1 positive lysosomes (n=20 dots). Each data point corresponds to the intensity of LAMP1 signal per field of view and size of lysosomes. (**B**) Confocal microscopy analysis of Cathepsin B (green) in cells indicating an increase in intensity of Cathepsin B protein in cells. Scale bar, 10um. Right graph: quantification of Cathepsin B signal relative to DAPI (n=4). Each data point corresponds to the intensity of Cathepsin B per field of view. (**C**) Western blot analysis of a lysosomal protease, Cathepsin B in cells indicating an increase in the unprocessed form of Cathepsin B on treatment with 27HC. (**D**) Western blot analysis of Cathepsin B in equal volumes of vesicle free media indicating an increase in Cathepsin B in media. (**E**) Quantitative PCR of Cathepsin B (CTSB) indicating an increase in mRNA (n=4). (**F**) Confocal microscopy analysis of autophagy marker LC3b (green) in cells indicating an increase in intensity of LC3b positive autophagosomes in cells. Scale bar, 10μm. Right graph: quantification of LC3b signal relative to DAPI (n=6). Each data point corresponds to the intensity of LC3b per field of view. Statistical analyses were performed using a Student’s t-test. ****P-value<0.0001, **P-value<0.01, *P-value<0.05. Data are presented as mean+/-SEM. Original confocal images were used for analysis and representative images depicted here were brightness adjusted.

An increase in the size of lysosomes has been suggested to occur as a result of lysosomal membrane permeabilization, where the integrity of the lysosomal membrane is disrupted, leading to a release of lysosomal components into the cytosol as well as their secretion into the extracellular space/media (47). If lysosomal membrane integrity is impaired by 27HC, we would expect an increase in lysosomal proteins secreted by the cells. Thus, we measured the expression of a lysosomal protease, Cathepsin B (CTSB), expecting that if lysosomes were disrupted there would be an increase in CTSB protein. Using IF, we found that 27HC significantly increased CTSB at the cellular level (**Fig. 3B**). CTSB is produced as pro-CTSB (unprocessed form) and gets cleaved in the lysosome to form active CTSB (processed form). We observed an increase in the unprocessed form of the protein by Western blot analysis (**Fig. 3C**). We also observed an increase in the amount of Cathepsin B protein in the conditioned media depleted of EVs (vesicle free media), indicating that the Cathepsin B is being leaked outside the lysosome and secreted outside the cells (**Fig. 3D**). Interestingly, the mRNA expression of *CTSB* was also increased by treatment of RAW cells with 27HC, suggesting that there may be feedback to replenish CTSB levels (**Fig. 3E**). Collectively, these data indicate that the lysosome is unable to efficiently cleave Cathepsin B protein and there is potential feedback to increase CTSB synthesis.

Lysosomes are also important for the process of autophagy, where autophagosomes transfer cytosolic contents to the lysosome by fusion to form autolysosome, and the contents get recycled within the cell (48). We observed that 27HC treatment leads to an increase in the amount of LC3b positive vesicles or autophagosomes within the cells (**Fig. 3F**), again suggesting lysosomal dysfunction is decreasing autophagosome processing. In summary, the data presented in **Fig. 3** strongly support our hypothesis that lysosomal function is being impaired by 27HC.

### 27-Hydroxycholesterol induces lysosomal dysfunction leading to an impaired acidification of lysosomes and increased EV secretion

To further study the effect of 27HC on lysosomes, we assessed its effect on lysosomal acidification. First, we assayed the number of acidic organelles in the cells using a LysoTracker probe, which is a cell-permeable fluorescent dye that stains acidic organelles such as the lysosome. The probe acts as a lysosomotrope; it diffuses across the plasma membrane and enters the acidic lysosomes, where it gets protonated and localized within the organelle. A decrease in LysoTracker signal was observed in cells treated with 27HC, indicating less acidic organelles (**Fig. 4A**). Next, we determined the pH of the cells using a LysoSensor probe which, is similar to the LysoTracker probe in that it accumulates in the lysosomes, but unlike the LysoTracker probe, the fluorescence intensity is dependent on the pH and increases with an increase in pH. There was an increase in intensity of LysoSensor signal in 27HC-treated cells, indicating an increase in lysosomal pH (i.e. loss of protons; **Fig. 4B**). These data indicate that lysosomes were failing to properly maintain their acidity, providing further support for 27HC disrupting lysosomal function.

**Figure 4:**
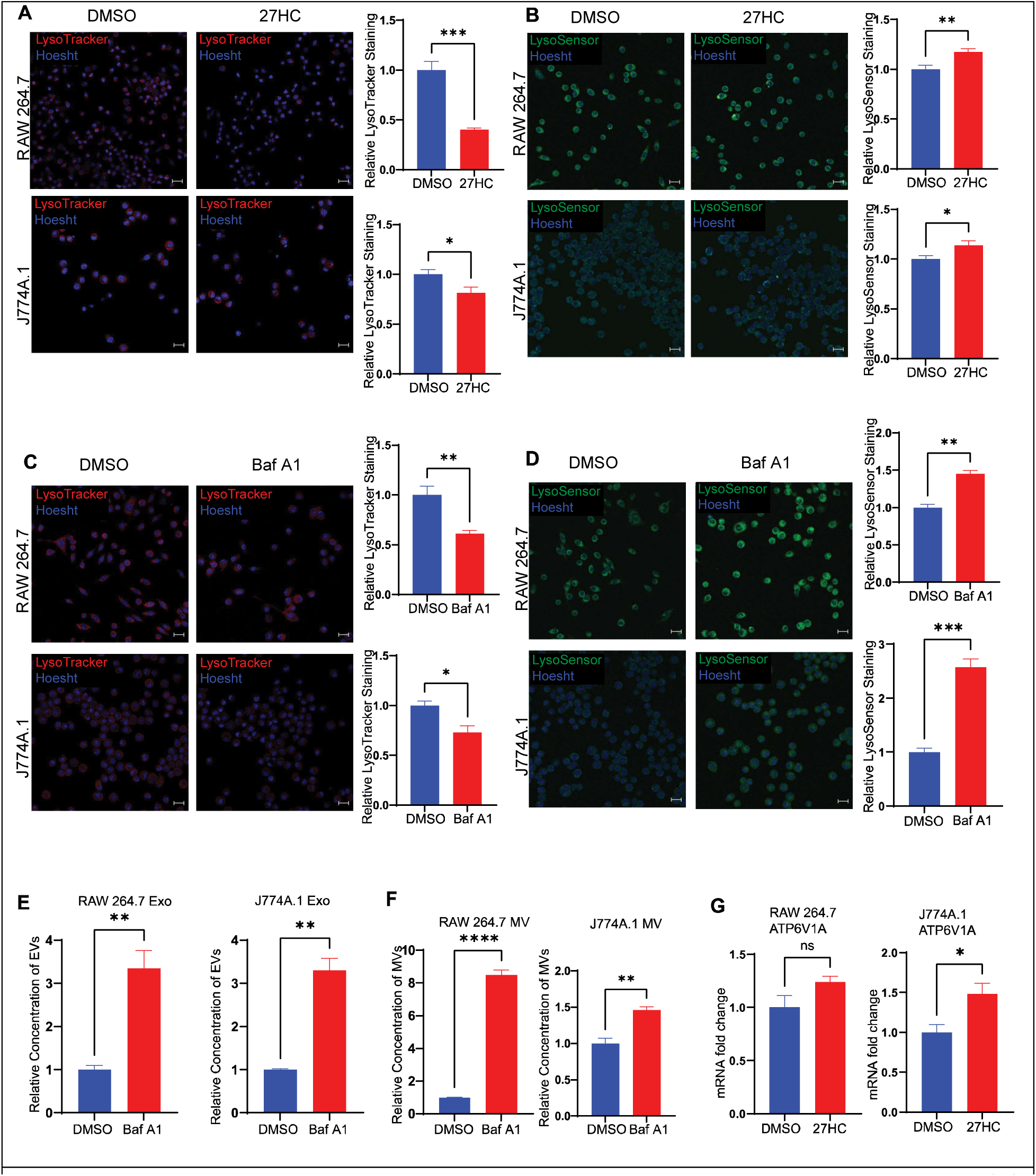
27HC impairs acidification of lysosomes which leads to increased EV secretion. (**A**) Confocal microscopy analysis of number of acidic organelles (lysosomes) using LysoTracker Deep Red showing a decrease in signal on treatment with 27HC, indicating decreased acidic organelles. Scale bar, 20µm. Right graph: quantification of LysoTracker signal relative to Hoesht, per field of view (n=4). (**B**) Confocal microscopy analysis of lysosomal pH using LysoSensor Green showing an increase in signal on treatment with 27HC, indicating an increase in pH. Scale bar, 20µm. Right graph: quantification of LysoSensor signal relative to Hoesht, per field of view (n=6). Treatment of cells with Bafilomycin A1, an inhibitor of the lysosomal V-ATPase pump ATP6V1A, lead to (**C**) a decrease in number of acidic organelles (n=4), (**D**) an increase in lysosomal pH (n=4), and (**E-F**) an increase in secretion of EVs (n=3). (**G**) Quantitative PCR of ATP6V1A indicating that 27HC does not change the levels of ATP6V1A at the mRNA level (n=4). Statistical analyses were performed using a Student’s t-test. ****P-value<0.0001, **P-value<0.01, *P-value<0.05. Data are presented as mean+/-SEM. Original confocal images were used for analysis and representative images depicted here were brightness adjusted.

The acidic lumen of lysosomes is in part maintained by ATP6V1A. ATP6V1A is a lysosomal V-ATPase, that functions by coupling the energy obtained from ATP hydrolysis to the transport of protons into the lysosomes (49). To study the effect of impaired lysosomal acidification on the rate of EV secretion, we treated cells with Bafilomycin A, a known inhibitor of ATP6V1A (50). As expected, treatment with Bafilomycin A led to a decreased number of acidic organelles and increased lysosomal pH (**Fig. 4C-D**). Importantly Bafilomycin A treatment increased the secretion of both small and large EVs (**Fig. 4E-F**), mirroring what we observed with 27HC treatment (**Fig. 1**). Our findings that Bafilomycin A treatment recapitulated the effects of 27HC lends strong support to our hypothesis that 27HC increases EV secretion by impairing lysosomal function, decreasing MVB degradation by lysosomes, and ultimately shifting them towards secretion as EVs.

Recent studies have shown that a decrease in ATP6V1A mRNA levels leads to an impairment of lysosomal acidification and an increase in EV secretion (15,43). If the pro-EV-secretory effects of 27HC were via ATP6V1A, we would expect 27HC to decrease its mRNA levels. However, when we examined the effect of 27HC on ATP6V1A mRNA levels, we observed no change in its expression in RAW 264.7 cells and an increase in expression in J774A.1 cells (**Fig. 4G**). Thus, 27HC likely acts upstream of, and not directly on the lysosome, to impair lysosomes.

### 27-Hydroxycholesterol increases intracellular oxidative stress and causes mitochondrial dysfunction

Our data thus far supports a mechanism whereby 27HC impairs lysosomal function, which leads to a decreased degradation of MVBs and an increase in the secretion of EVs. Our next step was to determine what contributes to this impairment of lysosomal function. Several studies have shown that elevated levels of reactive oxygen species (ROS) result in impaired lysosomal function (51,52). Interestingly, elevated ROS has also been found to increase EV secretion (53,54).

27HC treatment of cells increased levels of total cellular ROS (**Fig 5A**). This suggests that the mechanism by which 27HC disrupts lysosomes and subsequently increases EV secretion is via the initial production of ROS. Elevated levels of ROS can be a result of several factors including mitochondrial dysfunction (55–57). Further, altered mitochondrial function has been demonstrated to disrupt the structure and function of lysosomes in a ROS dependent manner (58–60). Therefore, we studied the effect of 27HC on mitochondrial function. 27HC treated cells had decreased total mitochondrial mass as measured using MitoTracker dye (**Fig. 5B**). TMRM staining indicated that there was also a decrease in mitochondrial membrane potential (**Fig. 5C**). Finally, we observed an increase specifically in mitochondrial ROS using the MitoSOX stain (**Fig. 5D**). Therefore, cellular exposure to 27HC leads to mitochondrial dysfunction and increased ROS; the ROS potentially disrupting lysosome function and thus increasing EV secretion.

**Figure 5:**
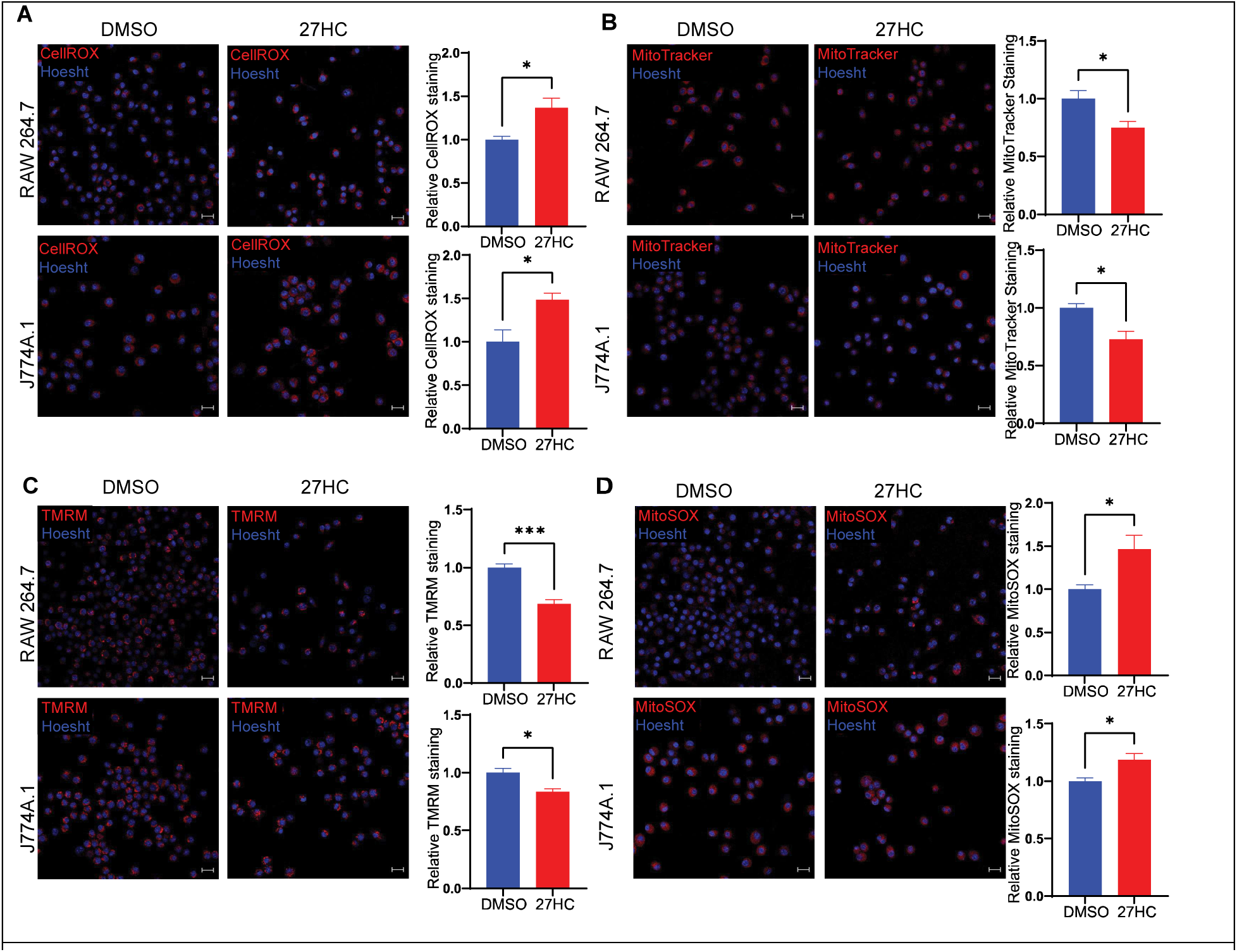
27HC increases intracellular oxidative stress and causes mitochondrial dysfunction. **(A)** Confocal Microscopy analysis of intracellular reactive oxygen species (ROS) using CellROX deep red indicating an increase in signal on treatment with 27HC, indicating increased levels of intracellular ROS. Scale bar, 20µm. Right graph: quantification of CellROX signal relative to Hoesht, per field of view (n=4). (**B**) Confocal Microscopy analysis of mitochondria using MitoTracker deep red indicating a decrease in signal on treatment with 27HC, indicating decreased mitochondrial mass. Scale bar, 20µm. Right graph: quantification of MitoTracker signal relative to Hoesht, per field of view (n=4). (**C**) Confocal Microscopy analysis of mitochondrial membrane potential using Tetramethyl Rhodamine Methyl Ester (TMRM) deep red indicating a decrease in signal on treatment with 27HC, indicating decreased mitochondrial membrane potential. Scale bar, 20µm. Right graph: quantification of TMRM signal relative to Hoesht, per field of view (n=4). (**D**) Confocal Microscopy analysis of mitochondrial ROS using MitoSOX deep red indicating an increase in signal on treatment with 27HC, indicating increase mitochondrial ROS production. Scale bar, 20µm. Right graph: quantification of MitoSOX signal per relative to Hoesht, per field of view (n=4). Statistical analyses were performed using a Student’s t-test. ****P-value<0.0001, ***P-value<0.001, **P-value<0.01, *P-value<0.05. Data are presented as mean+/-SEM. Original confocal images were used for analysis and representative images depicted here were brightness adjusted.

### 27-Hydroxycholesterol alters mitochondrial metabolism

Elevated mitochondrial ROS correlates with alterations in mitochondrial bioenergetics and overall cellular metabolic output (61). To study the simultaneous effects of 27HC on OXPHOS, TCA cycle and glycolysis within the cell, we used two techniques-Simultaneous Label-free Autofluorescence Multiharmonic (SLAM) Microscopy and real-time cell metabolic analysis (Seahorse Assays).

In a label-free manner, SLAM microscopy measures autofluorescence emitted by FAD and NAD(P)H in live cells. A decrease in the FAD:(FAD+NAD(P)H) ratio corresponds to an increase metabolic activity in the cells (62,63). We observed that 27HC treatment leads to an increase in the FAD:(FAD+NAD(P)H) ratio in RAW 264.7 cells (**Fig. 6A**). This would be reflective of a decrease in metabolic activity in the cells.

**Figure 6:**
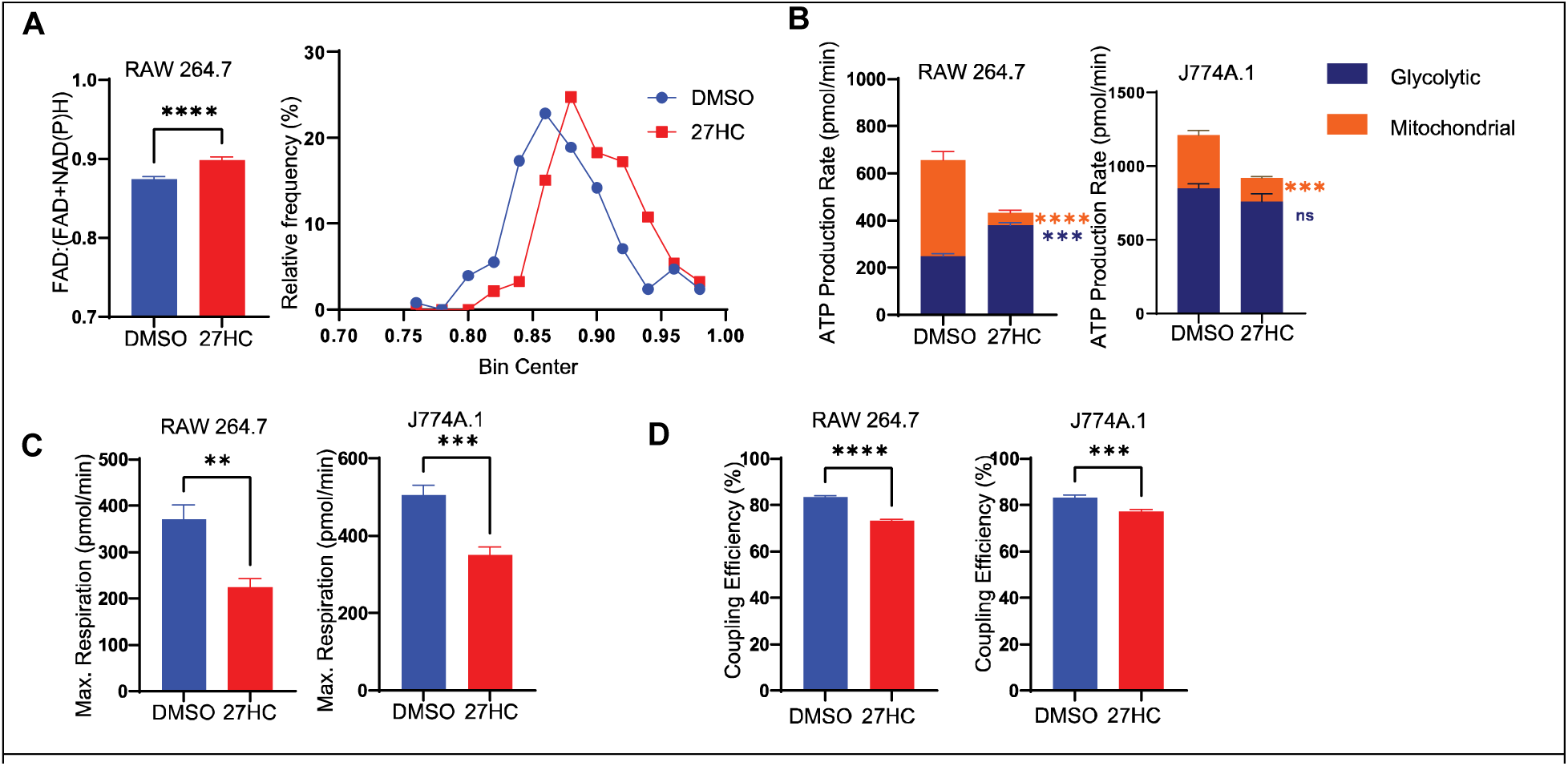
27HC alters mitochondrial metabolism in myeloid cells. (**A**) SLAM Microscopy in cells indicating an increase in FAD:(FAD+NAD(P)H) ratio in cells. **(B)** Seahorse ATP production Assay indicating a decrease in mitochondrial ATP production (n=6). **(C)** Seahorse Mito stress test indicating a decrease in levels of maximal respiration (n=6). **(D)** Seahorse Mito stress test showing decreased coupling efficiency in 27HC treated cells (n=6). Statistical analyses were performed using a Student’s t-test ****P-value<0.0001, ***P-value<0.001,**P-value<0.01. Data are presented as mean+/-SEM.

To more specifically assess metabolic flux, we measured oxygen consumption rate (OCR; an estimate of oxidative phosphorylation) and extracellular acidification rate (ECAR; an estimate of glycolytic activity), using these to calculate the proton efflux rate (PER). For both RAW 264.7 and J774A.1 cells, 27HC decreased OCR, with smaller changes observed in ECAR and PER (Seahorse ATP Rate Assay kit; **Supplementary Fig. 1A-C)**. Using these values mitochondrial versus glycolytic ATP production can be estimated. Notably, 27HC robustly decreased mitochondrial ATP production in both cell models, while increasing glycolytic ATP production in RAW 264.7 cells but not significantly changing glycolytic ATP production in J774A.1 (**Fig. 6B**). This was also reflected in the total ATP production rate in an independent experiment using the Seahorse Mito Stress kit (**Supplementary Fig. 1D-E**). Likewise, in an independent experiment using the Glycolytic Rate Seahorse Assay, a potential compensatory increase in estimated glycolysis was observed in RAW 264.7 cells, treated with 27HC, while no significant differences were found in J774A.1 cells (**Supplementary Fig. 1F-G**). 27HC lead to decreased maximum respiration rate and a reduced coupling efficiency, indicated by the Seahorse Mito Stress Test (**Fig. 6C-D**). Collectively, these data demonstrate that mitochondrial oxidative phosphorylation is robustly disrupted, suggesting that impaired mitochondria are responsible for the observed increases in ROS. Therefore, our data demonstrate a connection between mitochondrial function, lysosomal function and EV secretion; 27HC resulting in impaired mitochondrial function, increased ROS, disrupted lysosome function and thus a buildup of MVBs that are shunted towards secretion as EVs.

### A mitochondria-targeted antioxidant attenuates the 27HC mediated increase in EV secretion

Our evidence thus far implicates mitochondrial ROS as the upstream initiator of the effects of 27HC on EV biogenesis. Therefore, we co-treated cells with 27HC and a mitochondria-targeted antioxidant Mitoquinol (MitoQ). MitoQ is a potent antioxidant consisting of coenzyme Q10 linked to a lipophilic triphenylphosphonium cation to enable its accumulation in the mitochondria, which blocks the generation of ROS and prevents mitochondrial oxidative damage (64,65). We would thus expect MitoQ co-treatment to rescue the lysosomal damage. In support of our model, MitoQ was able to normalize LysoTracker signal after a 24-hour treatment with 27HC (**Fig. 7A**). Importantly, MitoQ was also able to attenuate the 27HC mediated increase in EV secretion over a 48-hour culture period (**Fig. 7B**). Therefore, we conclude that ROS is the upstream mediator of 27HC-stimulated EV secretion and use of an antioxidant can attenuate the increased EV secretion.

**Figure 7:**
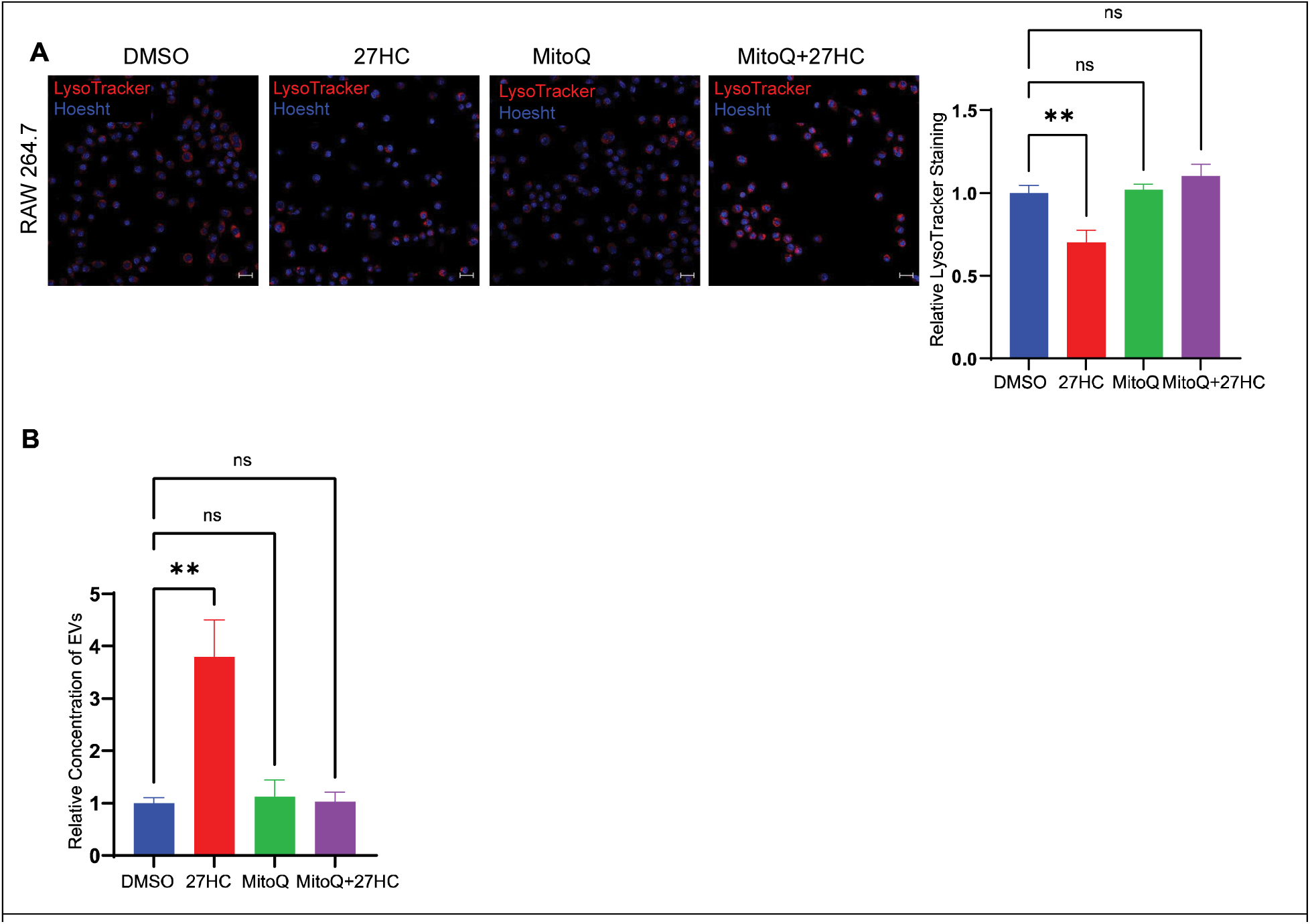
A mitochondria-targeted antioxidant attenuates the 27HC mediated increase in EV secretion. **(A)** Confocal microscopy analysis of number of acidic organelles (lysosomes) using LysoTracker Deep Red indicating that cotreatment with MitoQ normalizes the number of acidic organelles. Scale bar, 20µm. Right graph: quantification of LysoTracker signal relative to Hoesht, per field of view (n=4). **(B)** Nanoparticle tracking analysis of EVs obtained after 48 hours of treatment indicating that cotreatment with MitoQ attenuates the 27HC mediated increase in EV secretion (n=3). Statistical analyses were performed using one-way ANOVA with Dunnett’s multiple comparisons test. **P-value<0.01. Data are presented as mean+/-SEM. Original confocal images were used for analysis and representative images depicted here were brightness adjusted.

## DISCUSSION

EVs play an important role in maintaining regular cell to cell communication. EVs have also been strongly implicated as critical messengers in tumor progression and metastasis, in addition to other factors such as chemokines and growth factors (10,66). These EVs influence cancer progression by transporting molecules that can initiate several pathophysiological processes such as vascular leakiness, angiogenesis, and formation of the pre-metastatic niche (11,12). They also play a role in drug resistance, where the cells can directly efflux drugs via EVs (67). How or what factors regulate EV biogenesis and secretion is still poorly understood, although certain physiological stressors (e.g. hypoxia) and various metabolic factors have now been found to regulate EV secretion and/or cargo (3). The regulation of EV secretion is important to understand since it represents a promising strategy for the treatment of various diseases, including cancer.

Previous work has shown that treatment of myeloid immune cells with the cholesterol metabolite, 27HC, leads to an increase in the secretion of EVs, and these EVs promote tumor growth and metastasis (27). Given that there is an increase in the number of EVs and these EVs have pro-tumorigenic effects, understanding the mechanisms by which their secretion is increased would provide a unique opportunity to develop intervention strategies to prevent metastatic progression. We have demonstrated here that 27HC increases EV secretion by a cascade of alterations in cellular metabolism, characterized by impaired mitochondrial function, increased cellular oxidative stress and impaired lysosomal function.

The biogenesis of EVs of endosomal origin involves the maturation of endosomes to form multivesicular bodies (MVBs) containing intraluminal vesicles which either fuse with the lysosome and get degraded or fuse with the plasma membrane and get secreted as EVs (3). Therefore, the lysosomes determine the fate of MVBs; towards degradation and recycling, or towards secretion as EVs. Lysosomes help maintain cellular homeostasis by degrading and recycling cellular components. The impairment of lysosomes leads to the inability of the cell to recycle cellular components which are generated via the endosomal-multivesicular body pathway, leading to the secretion of these contents as EVs. We have observed here that 27HC induces lysosomal dysfunction as measured through increased lysosomal size and pH. This disrupts their inability to degrade multivesicular bodies and instead leads to the secretion of their contents as EVs. Several other groups have also demonstrated that lysosomes determine the fate of MVBs. One report has shown that hypoxia promotes EV secretion by impairing the lysosomal acidification (15). Another study found that a decrease in SIRT1 levels impairs lysosomal acidification and leads to secretion of EVs (43).

To further study what causes the observed impairment of lysosomal function, we studied the effect of 27HC on cellular ROS, since elevated levels of ROS results in impaired lysosomal function (51,52). We observed that 27HC increases levels of cellular ROS. A majority of the ROS produced in the cell is produced by mitochondria and dysfunctional mitochondria produce elevated levels of ROS (59,64). We have observed that 27HC impairs mitochondria and increases levels of mitochondrial ROS. Finally, since the upstream mediator for the increased EV secretion is the production of mitochondrial ROS, we co-treated the cells with a mitochondria-targeted antioxidant, Mitoquinol (MitoQ) and observed that MitoQ was able to rescue the lysosomes and attenuate the effects of 27HC on increased EV secretion.

Therefore, we have demonstrated that 27HC impairs mitochondrial function, which leads to elevated production of ROS. This elevated ROS in the cell disrupts the ability of lysosomes to degrade MVB cargo and leads to an increase in EV secretion. The use of antioxidants such as MitoQ can rescue these effects, by restoring lysosomal function and MVB degradation. Several *in vitro* and *in vivo* studies have demonstrated that MitoQ is protective against multiple diseases, including cancers (64,68–71). Therefore, potential intervention strategies to attenuate the 27HC induced increase in EVs could include the use of MitoQ or other antioxidants that can prevent oxidative damage.

Cancer progression involves crosstalk between various cells in the tumor microenvironment, including immune cells, stromal cells and fibroblasts. A vast majority of studies relating EVs and cancer progression involve the study of EVs derived from cancer cells and their influence on the functions of other cells in the tumor microenvironment (72–74). The role of immune cells in tumorigenesis is a double-edged sword. On one hand, immune cells help to enhance the anti-tumor immune response and act to eliminate cancer cells; while on the other, immune cells can be corrupted by cancer cells and become pro-tumorigenic. As a result, EVs from these cell types also play an important role in cancer progression. EVs from myeloid cells have been implicated in stimulating pro-metastatic properties, mediated by their interaction with other immune cells to form a suppressive tumor microenvironment (11,75,76). In the context of elevated cholesterol and breast cancer, we have seen that EVs derived from 27HC treated neutrophils promote breast cancer progression (27). Other studies involving immune cell derived EVs have revealed that EVs containing miR-501-3p derived from tumor associated macrophages (TAMs) are involved in cell migration and invasion leading to the progression of pancreatic cancer and the suppression of this miRNA in cells can inhibit this progression (77). TAMs can also promote invasion of breast cancer cells by transfer of miR-223 via the Mef2c-β-catenin pathway (78). Macrophage derived EVs can reverse breast cancer cell quiescence and induce proliferation, leading to re-emergence from dormancy (79).

Altered cholesterol metabolism has been strongly associated with the progression of various diseases. Interestingly, several miRNAs have been implicated in the regulation of cholesterol homeostasis (80). Furthermore, cholesterol derivatives other than 27HC have also been found to alter EV secretion with altered cargo. For example, dendrogenin A, was found to stimulate cancer cells to secrete EVs carrying altered cargo that increased dendritic cell maturation and Th1 T lymphocyte polarization (81). When considering our work along with other published studies, it becomes clear that EVs play an important role in the progression of cancer, but that more insights into their regulation are required for clinical translation of this knowledge.

Understanding the mechanisms that cells use to modulate EV secretion has been a long-standing goal in understanding normal cellular physiology, as well as changes in diseases such as cancer. Overall, a more comprehensive understanding of the regulation of EV secretion will help open avenues in identifying targets that can be modulated to rescue the increase in EV secretion. Our work has demonstrated the effect of elevated levels of a cholesterol metabolite, 27HC, in increasing EV secretion. This knowledge will help identify targets or metabolites in patients as prognostic markers for diseases such as cancer and other metabolic diseases and act as druggable targets as a therapeutic strategy.

## Funding

Department of Defense Era of Hope Scholar Award BC200206/W81XWH-20-BCRP-EOHS (ERN)

National Institutes of Health grant R01 CA234025 (ERN)

National Institutes of Health grant T32 GM136629 (HEVG)

National Institutes of Health grant T32 ES007326 (ATN)

National Institutes of Health grant P41 EB031772 (SAB)

Cancer Scholars for Translational and Applied Research (C*STAR) Program sponsored by the Cancer Center at Illinois and the Carle Cancer Center CST EP012023 (HEVG)

Postdoctoral Fellows Program at the Beckman Institute for Advanced Science and Technology (NK)

School of Molecular and Cellular Biology Summer Undergraduate Research Fellowship (HK)

## Acknowledgements

SThe Cancer Center at Illinois Tumor Engineering and Phenotyping Shared Resource (TEP) provided certain cell lines, routine mycoplasma testing and Seahorse XF assays (directed and assisted by Hui Xu and Huimin Zhang). TEM was performed by Catherine L Wallace at the Beckman Institute Imaging Technology Group Microscopy Suite.

## Supplementary Figures

**Supplementary Figure 1:**
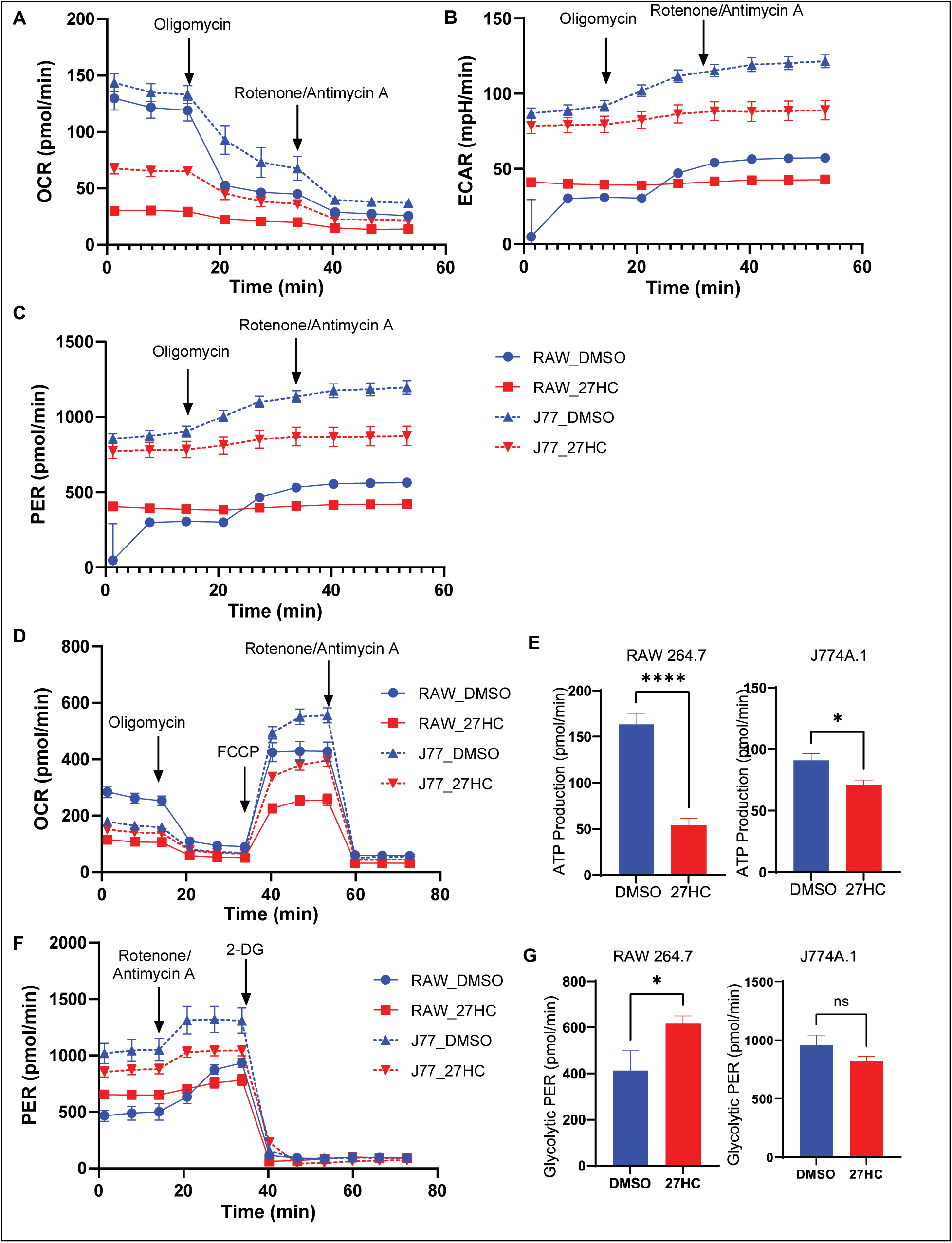
**(A)** Seahorse ATP Production Assay indicating oxygen consumption rate (OCR), **(B)** extracellular acidification rate (ECAR) and **(C)** PER rates (n=6). **(D)** Seahorse Mito Stress Test indicating OCR and **(E)** Mitochondrial ATP production rate (n=6). **(F)** Seahorse Glycolysis Stress Test indicating PER and **(G)** glycolytic PER (n=5/6). Statistical analyses were performed using a Student’s t-test ****P-value<0.0001, *P-value<0.05. Data are presented as mean+/-SEM.

