## Supplementary Figure 1 for "Cholesterol Metabolite 27-Hydroxycholesterol Enhances the Secretion of Cancer Promoting Extracellular Vesicles by a Mitochondrial ROS-Induced Impairment of Lysosomal Function"

### Supplementary Figures

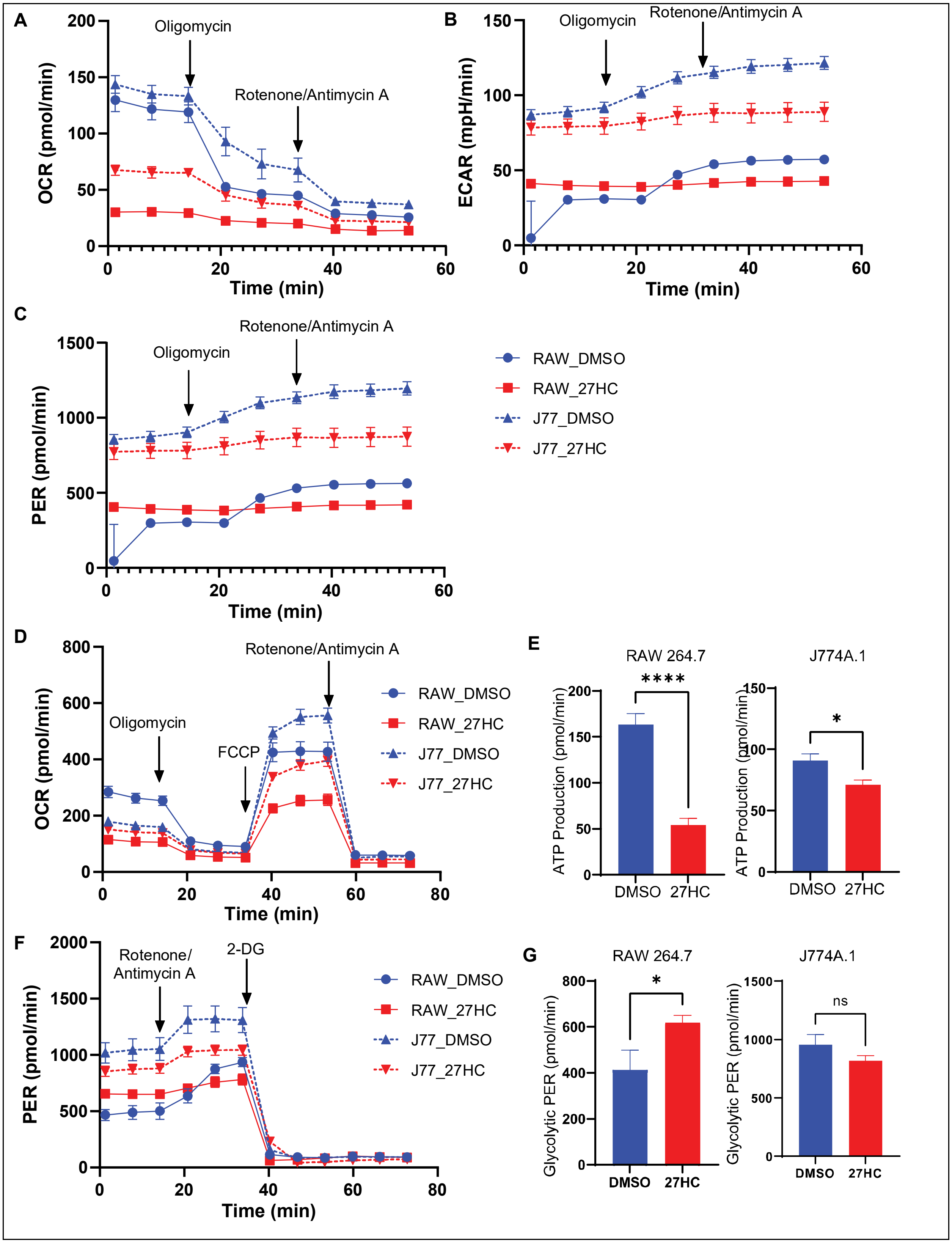
